# ATP induced conformational change of axonemal outer dynein arms studied by cryo-electron tomography

**DOI:** 10.1101/2022.05.02.490280

**Authors:** Noemi Zimmermann, Akira Noga, Jagan Mohan Obbineni, Takashi Ishikawa

## Abstract

Axonemal dyneins in the outer dynein arm (ODA) generate force for ciliary beating. We analyzed three states of ODA during the power stroke cycle using in situ cryo-electron tomography, subtomogram averaging and classification. These states of force generation depict the pre-power stroke, post-power stroke conformations and an intermediate state. Comparison of these conformations to published in vitro atomic structures of cytoplasmic dynein, ODA and Shulin-ODA complex showed that the orientation and position of the dynein head and linker differs. Our analysis shows that all dynein linkers in the in the absence of ATP interact with AAA3/AAA4, indicating interaction to the adjacent B-tubule direct dynein orientation. For the pre-power stroke conformation, we found changes of the tail anchored on the A-tubule. We built pseudo-atomic models from high-resolution structures to generate a best fitting atomic model to the in-situ pre- and post-power stroke ODA, thereby showing that the Shulin-ODA display similar conformation as the active pre-power stroke ODA conformation in the axoneme.

## Introduction

Motile cilia are rod-like extensions protruding out from the cell body. Their function is to provide movement, either to the organism itself or to its near environment. They exist from unicellular eukaryotes such as *Chlamydomonas* and *Tetrahymena* to human. In human, motile cilia function in various tissues, such as the lung, kidney, brain and reproductive organs and their defect causes ciliopathies, such as primary ciliary dyskinesia (PCD) (Hildebrandt et al., 2011). The beating motion of cilia takes place at the part extending out of the cell body, called axoneme. In most species, the axoneme forms a “9+2” structure, in which nine microtubule doublets (MTD) surround two singlet microtubules. The MTD is made of a complete cylindrical microtubule (MT) with 13 protofilaments (A-tubule) and an incomplete B-tubule with 10 protofilaments (Ma et al., 2019). Until now more than 400 proteins are known as components of motile cilia (Pazour et al., 2005).

The driving force of ciliary motion are axonemal dyneins. They are anchored on a microtubule doublet and generate force against the adjacent doublet by hydrolyzing ATP. In most species, axonemal dyneins compose two complexes: outer and inner dynein arms, which make periodic arrays with 24nm and 96nm periodicity, respectively. The outer dynein arm (ODA) has either two or three heavy chains (HC), stacking on the doublet (Goodenough, 1982; Ishikawa et al., 2007; Nicastro et al., 2006; Rao et al., 2021; Walton et al., 2021), whereas eight heavy chains make an array in the inner dynein arm (IDA). Functionally ODA is responsible for force generation, whereas IDA is responsible for regulatory functions (Kamiya, 2002). Each heavy chain of the axonemal dynein consists of an N-terminal tail domain followed by the motor domain, including six AAA motifs (named AAA1 – AAA6), with a microtubule binding domain (MTBD) connected to AAA4 by a coiled-coil stalk, similarly to cytoplasmic dynein, its homologue. The α-helix rich linker region is between the tail and AAA1 (Carter et al., 2011; Kon et al., 2011). Additionally to the HC, there are a number of light and intermediate chains present in the ODA and IDA. The tail complex of the ODA is formed by the N-terminal tails of the HCs (Kubo et al., 2021; Mali et al., 2021; Rao et al., 2021; Walton et al., 2021). For *Chlamydomonas* the β and γ tails form the scaffold for the two intermediate (IC1 and IC2) and multiple light chains (IC and LC). Most of the LC form the light chain tower (LC-tower), which sits on the γ tail. Only LC1 and LC4 are associated with the γ tail at the outside of the LC tower. LC3 and the α HC via its kelch domain bind to the β tail. The α HC docks on the β tail at helical bundle 6 (Rao et al., 2021). The N-terminal end of the γ and β tail form the N-terminal dimerization domain (NDD). This tail complex is anchored near the NDD on A-tubule by DC3 as well as DC1 and DC2.

Both axonemal and cytoplasmic dyneins make power stroke (PS) with respect to the microtubule, releasing γ phosphate after ATP hydrolysis (Johnson, 1985, 1983). Single particle cryo-EM (SPA) study of the motor domain from axonemal dynein demonstrated the swing of the linker during the power stroke (Roberts et al., 2009). The same conformational change of the motor domain from cytoplasmic dynein has been studied at atomic resolution (Carter et al., 2011; Schmidt et al., 2015). The linker interacts with the interface of AAA4 in the post-PS conformation and AAA2 in the pre-PS conformation (Carter et al., 2011; Kon et al., 2012; Schmidt et al., 2015), which is consistent with single particle cryo-EM work of isolated axonemal dynein (Roberts et al., 2009) and cryo-EM/ET work of ODA in the axoneme (Kubo et al., 2021; Lin and Nicastro, 2018; Ueno et al., 2014; Walton et al., 2021) at intermediate resolution. Recent high resolution SPA has shown that there are two different linker conformations present in the axonemal ODA of *Tetrahymena* (Kubo et al., 2021; Rao et al., 2021)for the post-PS structure, the so called Post-1 and Post-2 conformation. While in the Post-1 conformation, the linker goes over AAA4, in the Post-2 conformation the linker interacts with the interface of AAA3 and AAA4. The Post-2 conformation was this far only observed for DHY4 and DHY5 (β and α HC analog in *Chlamydomonas*).

A recent work revealed that in *Tetrahymena* during intraflagellar transport (IFT) of the ODA, the ODA is in a closed conformation, where the DHY3 (γ HC analog) is folded onto the other two HC (Mali et al., 2021). This conformation is held together and inhibited by Shulin. In this Shulin-ODA complex the dynein linker adapt a pre-PS conformation. The stalkes are curved and interact with each other close to their MTBDs. Once Shulin falls off in the axoneme, the ODA is no longer inhibited, opens up and binds to the A-tubule. For this the DHY3 undergoes a 90° rotation together with DIC2 (IC1 analog) and the LC tower.

The interaction between the MTBD and the B-tubule has been studied by cryo-EM. MTBD binds on a protofilament of the B-tubule, between α- and β-tubulins, under the strong binding condition, corresponding to the post-PS (Carter et al., 2008; Redwine et al., 2012),but change its binding angle at the pre-PS (Redwine et al., 2012). The stalk, protruding from AAA4, tilts toward the proximal direction (minus end of MT) both at pre- and post-PS condition (Lin et al., 2014; Movassagh et al., 2010; Ueno et al., 2014, 2008), indicating that sliding motion between two adjacent doublets occur by dynein, which is anchored on the A-tubule, dragging the adjacent B-tubule (winch model) (Movassagh et al., 2010; Ueno et al., 2008), rather than rotating to push the B-tubule (rotation model). What is not known is the precise conformational change of the axonemal dynein head during PS, due to lack of resolution to image dyneins generating force against MT.

Single particle cryo-EM successfully revealed 3D conformational change of the motor domain of isolated cytoplasmic and axonemal dyneins. However, this method so far missed out on information of the ODA in cilia with and without ATP. This information knowledge is necessary to explain how ODA generate force in cilia and to explain how ciliary beating occurs. For that we tried to visualize the entire ODA structure in cilia with and without ATP by cryo-electron tomography (cryo-ET). To describe the structural change, we fitted published atomic structure of ODA, either anchored on the A-tubule but isolated from the B-tubule (Kubo et al., 2021; Walton et al., 2021) or connected to the B-tubule, but not anchored on the A-tubule (Rao et al., 2021) to our structure reconstructed by cryo-ET. First we described how ODA structure in the axoneme differs from those structures analyzed by single particle cryo-EM and built pseudo-atomic model of post-PS ODA structure. Next, we used molecular fitting to build a model of pre-PS ODA structure. Based on these models, we discuss mechanism of force generation of ODA and interaction with adjacent B-tubule at pseudo-atomic resolution.

## RESULTS

Relion classification of subtomograms from *Chlamydomonas* axoneme in the presence of ATP led to three major classes. One of these classes represents the MTD-1, which was previously shown to lack ODA (Bui et al., 2012). From the two remaining classes, one represents the pre-PS conformation as shown later. The other is likely representing a conformation after post-PS and before pre-PS, judging from the position of the dynein head more proximal than in the post-PS conformation and more distally than the pre-PS conformation. Therefore, we called this conformation intermediate conformation. No post-PS like conformation was found in these subaverages. Therefore, we obtained an apo state, depicting the post-PS conformation from a nucleotide free dataset. The resolution based on the FSC is 30Å for the post-PS conformation and 36Å for the pre and intermediate conformation (Supplementary Figure 1C).

### Post-PS structure and Fitting of the atomic models

Recently several atomic structures of ODA in the post-PS were solved by single particle cryo-EM (PDB-7KZM, 7MOQ, 7K58 and 7K5B) (the list of PDB files in the supplementary table). These structures are from specimen where the ODAs are either anchored on the A-tubule or isolated and reconstituted on the adjacent B-tubule. All these structures are without additional nucleotides. We attempted to model pseudo-atomic structures of ODA by fitting atomic models to our cryo-ET subtomogram averaging map without additional nucleotide (Figure 1).

**Figure 1.**
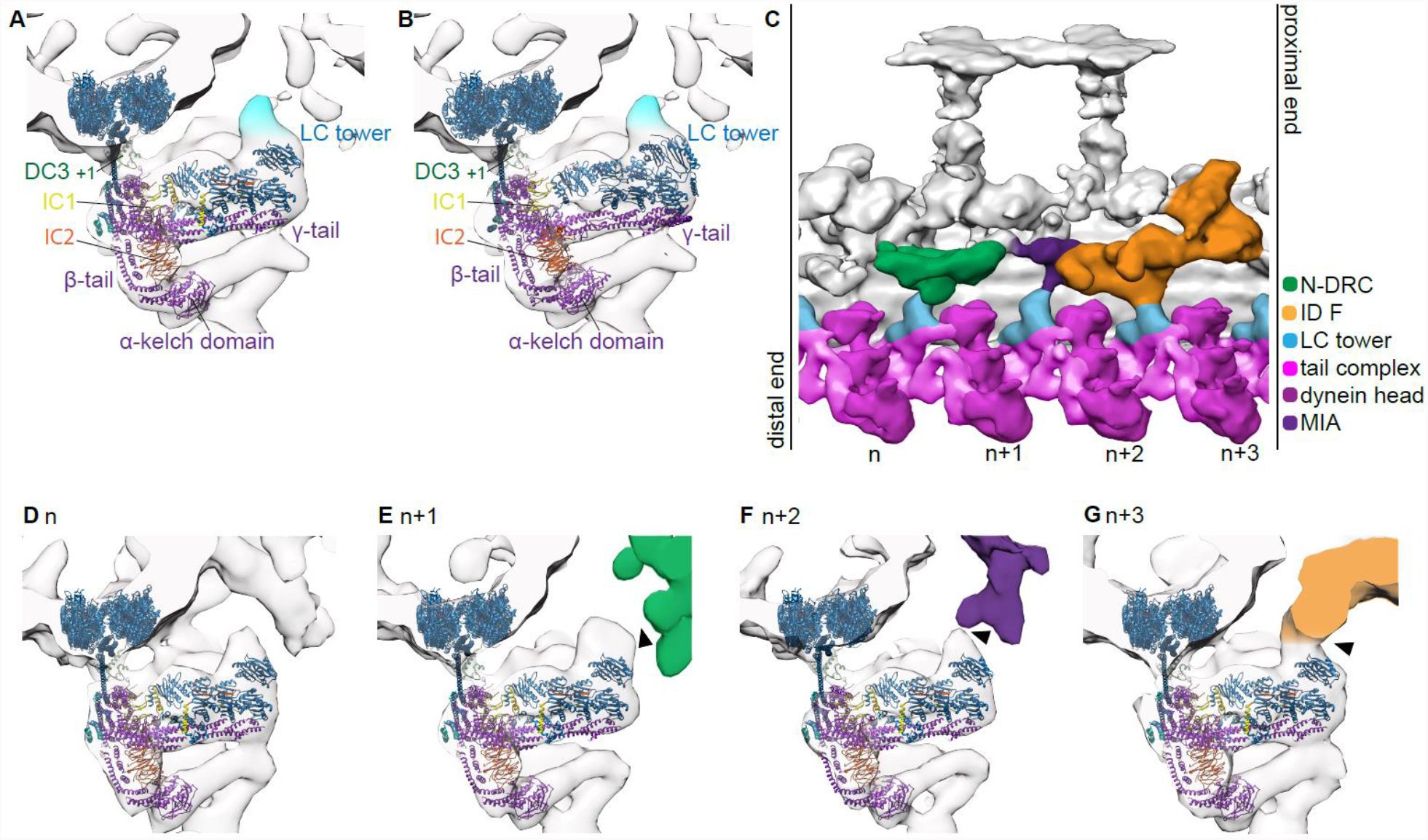
Tail ODA structure by subtomogram averaging from cryo-ET of Chlamydomonas cilia. A and B) LC tower of 24nm subtomogram average of Chlamydomonas, with the atomic model PDB-7KZM (A) and PDB-7MOQ (B) fitted. Fitting was done to maximize the cross-correlation between the whole ODA structure and the tomographic map. An unassigned density above the LC tower is highlighted in light blue. C) 96nm unit from post-PS Chlamydomonas by subtomogram averaging. D-G) Each LC tower of the 96nm average viewed from the end-on view. The LC tower connects once every 96nm to the IC/LC complex of inner dynein f (G, “n+3”) and to the MIA complex (F, “n+2”). The LC tower also comes close to the N-DRC (green) but does not seem to form a strong interaction with it (E, “n+1”). All outer dynein to inner dynein connections are indicated by black arrows.

In all four atomic models, the LC tower, fitted as a rigid body, does not completely fill out our tomographic density: in each 24nm repetitive unit there is an additional density above the LC tower (Figure 1A and B). This density comes close but does not connect to the N-DRC (“n+1” in Figure 1C,E). On the contrary, the density connects more strongly to the IC/LC of inner dynein f and the MIA complex (Yamamoto et al., 2013) (“n+2” and “n+3” in Figure 1C,F,G).

Both models (PDB-7KZM (Walton et al., 2021) and PBD-7MOQ (Kubo et al., 2021)), in which ODA is anchored on the A-tubule, were first fitted based on the DC3 and the PF. With this alignment, the γ HC of the PDB-7MOQ model is slightly (approx. 2nm) more distal than in the PDB-7KZM. The 7KZM fits better to the in-situ structure (Supplementary Figure 2A,C; Supplementary Movies 1 and 2). The buttress/stalk and linker position (AAA4) for the γ dynein are very similar between the two models, pointing a 1-2 nm more distal than the tomographic map suggests. Individual fitting of the γ head of PDB-7KZM results in a rotation and translation of the head with respect to the tail, increasing the fit between the map and the model (Figure 2A-B,C; Supplementary Movies 1). However, the position of the linker is less in agreement with the tomographic map (Supplementary Figure 2A). If this ever so slight head rotation would pull the C-terminal end of the tail closer to the A-tubule, which is indeed suggested by the tomographic map, the γ dynein linker would result in being in a Post-2 conformation, where the linker spans over AAA3/AAA4 (Kubo et al., 2021 and Rao et al., 2021). A pseudo-atomic Post2 conformation of the γ linker (Figure 2F) highlights that in this conformation the stalks, head, linker and C-terminal tail would fit into the tomographic map (Supplementary Movies 3).

**Figure 2.**
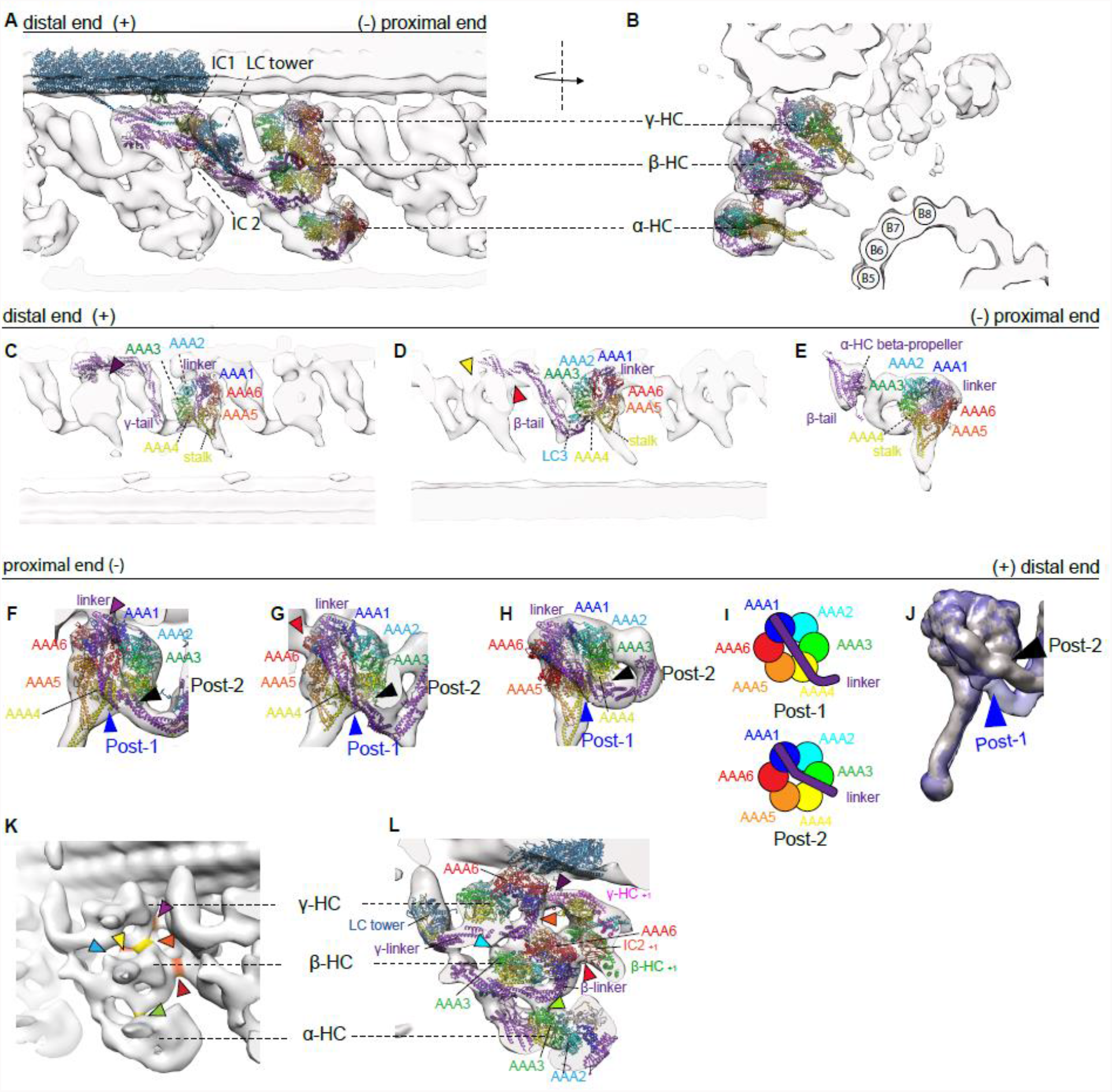
Comparison of atomic models single particle analysis post-PS ODA reconstituted from the A-tubule and our cyro-ET subtomogram averaging map. A and B) Composition of the best rigid body fitting atomic model structures into our tomographic map. (LC, IC, γ and α HC from PDB-7KZM (Walton et al., 2021), β HC from PDB-7MOQ (Kubo et al., 2021)). IC, LC and the γ tail were all together fitted as a rigid body. The γ (C) and α heads (E) were fitted separately from the rest of the ODA structure as rigid bodies. D) B HC of PDB-7MOQ fitted as one rigid body into the tomographic map. The SPA structure fits perfectly into the map. E) A 2 nm shift was enough to fit the α head into the tomographic map. Although there is still a stalk which points upwards in the atomic model the tomographic map suggest a straight conformation. F-H) The linker region of the pseudo atomic post-PS model fitted in the post-PS map. All three linkers adopt a Post-2 (black arrowhead) rather than a Post-1 conformation (blue arrowhead). The red and purple arrowheads indicate the head to tail interactions to the proximal ODA. I) Schematic representation of Post-1 and Post-2 linker conformations (Rao et al., 2021). J) 20Å map calculated from a Post-1 (blue) and Post-2 (black) conformation dynein overlaid. K) Intra (orange) and inter-ODA (yellow) connections. L) Pseudo-atomic post-PS model fitted into the same view of K to indicate which proteins are involved in the inter- and intra-ODA connections. The following interactions are marked by arrow heads: purple-γ AAA6 to proximal γ tail, orange and light blue – γ linker to β head, yellow-β head to proximal NND, red-β head to proximal IC2, green – β linker to α head.

More discrepancy can be found between the β HC of the two models. Whereby the structure of Kubo et al. overlays perfectly with our tomographic map (Figure 2A-B,D). Their proposed linker and LC3 position also aligns with our tomographic map (Figure 2G). The linker position for the post-PS conformation matches previous findings in Kubo et al. where they propose that the linker spans over AAA3/AAA4 (Post-2). Walton et al. found that the stalk of the β HC points distally in the majority of their particles. So far, no such structure was observed in our subtomogram averages (Supplementary Figure 2D).

When we fitted models with the third HC, either α of *Chlamydomonas* or DHY5/γ of *Tetrahymena* (PDB-7KZM, 7K58 and 7K5B), we found an additional density next to the α dynein head visible in the tomographic map (Supplementary Figure 2G-I), suggesting that in the in situ structure there is an additional protein present next to AAA2/AAA3, which probably fallen off or smeared out due to flexibility during single particle analysis. It might be the density of one of two known binding partners of the α HC are Lis1 and LC5 (King, 2016). In *Chlamydomonas* Lis1 binds as a monomer (Rompolas et al., 2012, p. 1). Using one unit of the dimeric *Homo sapiens* Lis1 (PDB-5VLJ (Htet et al., 2020, p. 1)) and fitting it into our density allowed us to assess its likeability. The volume of Lis1 corresponds to the volume of the unassigned density. Lis1 or any other binding partner would bind to the interface between AAA2 and AAA3 of the α dynein as well as the Ig-Fln of the α tail (Supplementary Figure 2H,I). The other discrepancies between the atomic models and the tomographic map of the α HC are most likely due to displacement. The α dynein head of 7kmq appears too proximal and close to the adjacent B-tubule (Supplementary Figure 2G). Individually fitting the α head of PDB-7KZM results in a good fit between the atomic model and the subtomogram average (Figure 2 A-B,E). This individual fitting changes the distance between the kelch-like β propeller and the dynein head, moving them closer together. So that the distance between the kelch-like β propeller and the head structure is larger in single particle analysis than in cryoET (Supplementary Figure 2G).

Inter and intra ODA interactions appear to be as described by Walton et al. and Kubo et al. for the γ and β HC. The γ HC connects to the proximal γ tail (helical bundle 3) close to the DC3 and via the linker to the β HC within the same ODA (Figure 2C,K,L). The β HC connects to the proximal tail structure at the region of IC2 (Figure 2D,G,K,L) and also to the proximal NDD (Figure 2B,K,L). An additional connection is observed from the tomographic maps between the AAA3 α HC and the β linker (Figure 2K,L).

The structures generated by reconstitution of ODA on the MTD (MTBS1 (PDB-7K58) and MTBS2 (PDB-7K5B) (Rao et al., 2021)) were also fitted to our tomographic map via rigid body fitting (Figure 3A-D) (Supplementary Movies 4). Neither of the two structures represented our conformation of the ODA. As mentioned above the tomographic map suggests that the γ linker is in a Post-2 conformation, which was not observed in neither MTBS1 nor MTBS2.

**Figure 3.**
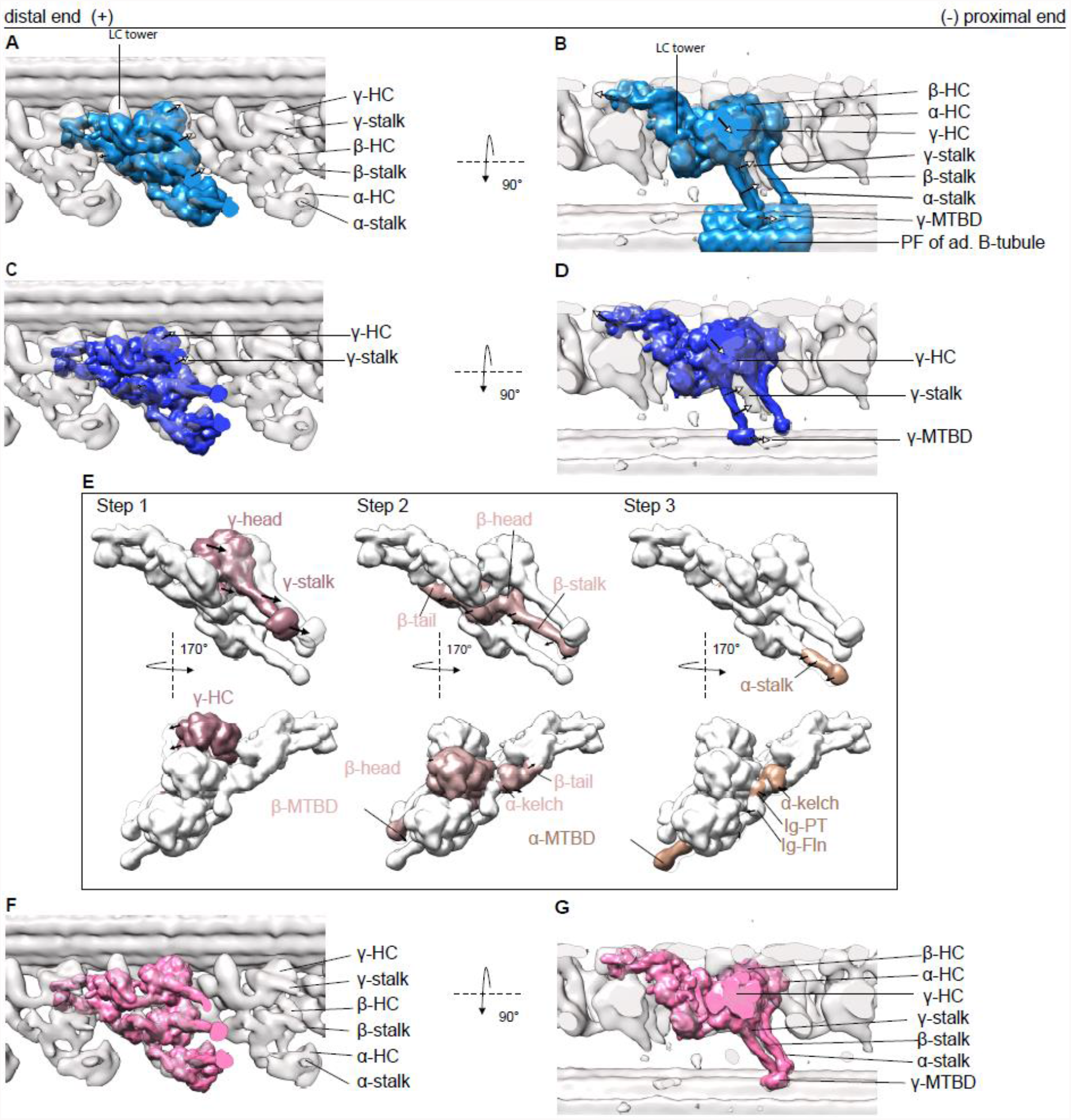
20Å resolution map calculated from the MTBS1 and MTBS2 (PDB-7K5B and PDB-7K58) model and fitted to our tomographic map to highlight the difference of stalk B-tubule binding between the in situ and in vitro structures. A and B) Rigid body fit of the whole MTBS1 map (Walton et al., 2021). There is discrepancy at the dynein heads, including the stalks and the MTBDs (indicated by the arrows) and the tomographic map. C and D) Rigid body fit of the whole MTBS2, there biggest difference between the map and the model is in the γ dynein head, stalk and its MTBD. There are also some smaller discrepancies for the β and α HC. E) Steps necessary to fit the MTBS2 model to the tomographic map. Step1: The γ head needs to shift. Changing the linker position from a Post-1 conformation to a Post-2 conformation. Step 2: The β HC, from helical bundle 5 to the β dynein head, shifts slightly. The two binding partners LC3 and the α kelch domain move with the β tail. Step 3: The stalk and MTBD of the α HC shifts. The α kelch domain moves closer to the N-terminal end off the β tail. F and G) Real space refined model after the three steps.

Since all the three linkers in situ appear to be in a Post-2 conformation, MTBS2 (Figure 3C-D), which has the β and α homolog linker in a Post-2 conformation, fits better than MTBS1 (Figure 3A-B). To optimize the fitting of the MTBS2 to the cryo-ET map we performed rigid body fitting and real space refinement of individual parts of the structure by Coot (Figure 3E). This optimization included three steps. Step 1 includes the translation and rotation of the γ dynein head including the MTBD. This shifts the MTBD by 8nm, which corresponds to one step on the adjacent B-tubule. The linker of the γ HC changes from a Post-1 conformation to a Post-2 conformation. Step 2 includes a 1nm backwards rotation of the β HC form helical bundle 5 up to the β dynein head. In addition to the β HC, this also moves LC3 and the α kelch domain so that they fit better into the tomographic map. Step 3 focuses on the α HC, with the kelch domain moving 1nm closer to helical bundle 5 of the β HC. In addition, the stalk of the α HC adapts to a more straight conformation by 1nm distal movement of the MTBD. The resulting MTBS2’ map is shown in figure 3F-G. By this remodeling the cross correlation score between the 20Å filtered MTBS2 and MTBS2’ structures and the cryo-ET map increased from 0.6797 to 0.8402.

### Pre- and intermediate-structures

In the presence of ATP, RELION 3D classification detected two major conformations of ODA in the wild type *Chlamydomonas* cilia. One resembles the previously known pre-PS state since (1) the positions of the heads are ∼8nm proximal to the post-PS structure, similarly to the ODA structure with ADP and vanadate (Movassagh et al.) and (2) the orientation of the linker is the same as cytoplasmic dynein with ADP and vanadate (Schmidt et al.). The other structure, which we define as an intermediate conformation and shares a lot of pre-PS like features, such as dynein head position closer to pre- than post-PS and linkers in a pre-PS like conformation (Figure 4D-I). Alignment of the different tomographic maps was done based on the RS, which led the DC3 to overlap (Figure 4A-B). The dynein head moves from their distal post-PS conformation, via their intermediate position to their most proximal pre-PS conformation (Figure 4D-F). With this movement the linker changes from a straight post-PS conformation to a 90°-bent pre-PS conformation in the intermediate and pre-PS structure (more detail below)(Figure 4G-H). For the pre-PS the stalks are more strongly tilted towards the adjacent B-tubule than for the intermediate structure (Figure 4D-E). Although classification was done based on only the central ODA, there are more global changes found outside of the ODA. By overlaying (Figure 4J) it becomes apparent that the distance and angle between the two adjacent MTDs changed, whereby the angle between two adjacent MTDs in the pre-PS conformation is wider (approx. 40°) than for the intermediate conformation (approx. 34°) (Figure 4J). In addition to this slight rotation of the adjacent MTD, in the intermediate conformation the distance between the ODA and MTD is approx. 2nm longer compared to the pre-PS conformation (movement indicated in Figure 4J).

**Figure 4.**
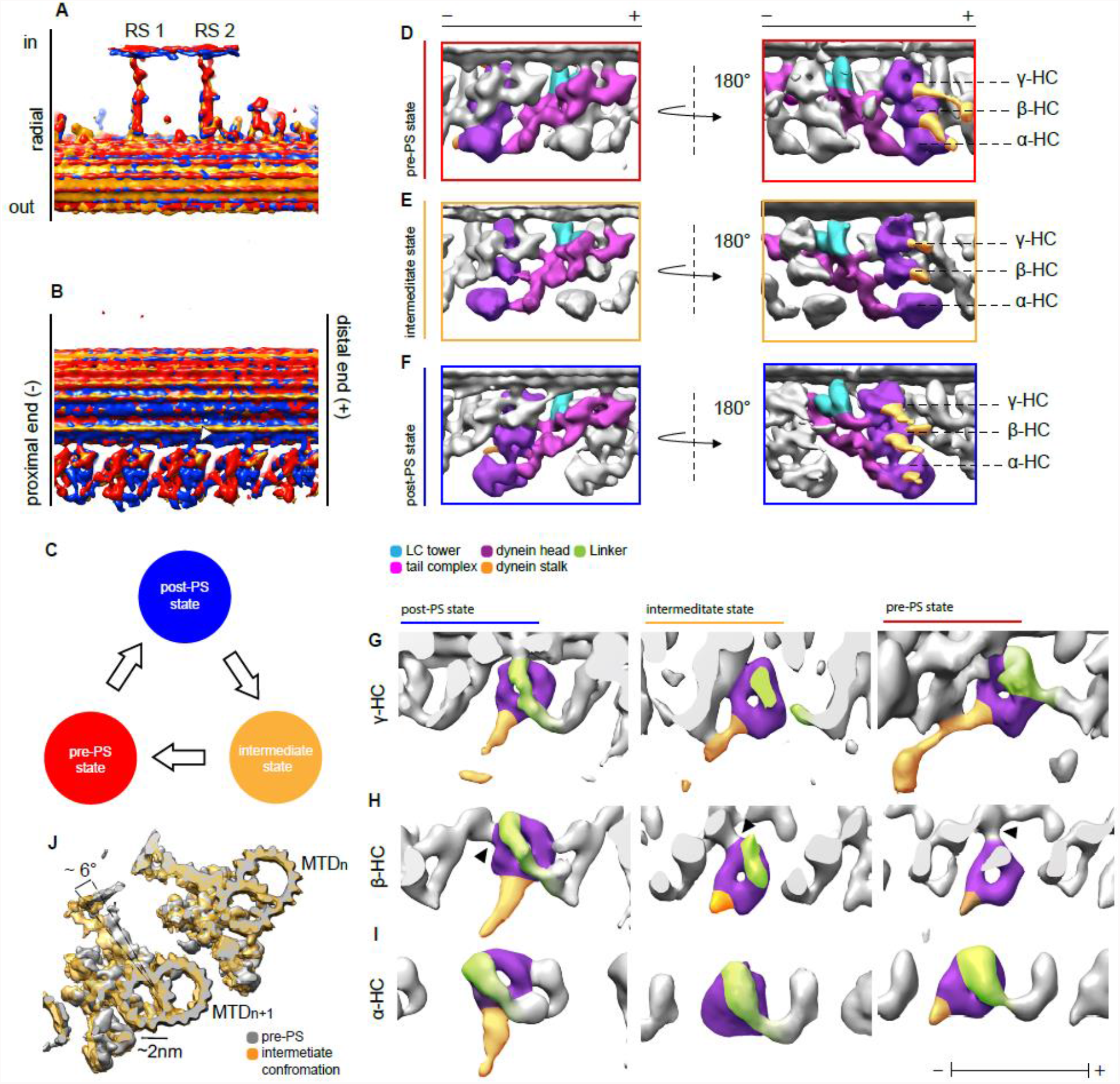
Structural changes of ODA between different states. A) Overlay of the 96nm array of the three states (post-PS, pre-PS and intermediate state). The structures are aligned based on the RS. With this the DC3 (indicated by white arrowhead) are also aligned. C) Model of the 3 states. The intermediate state is after the post and before the pre-PS states. The dynein heads move toward the proximal direction in the intermediate (E) and pre-PS state (D) compared to the post-PS state (D). Also the LC tower moves in the pre-PS state compared to the post-PS. γ (G), β (H) and α (I) HC in the post-PS, intermediate and pre-PS conformation. Linker position for all the HC and states are indicated in green, while the stalk and the AAA-ring are shown in yellow and purple, respectively. The black arrowhead in H indicates the interaction of the β dynein head to the proximally located IC2, which is consistent in all three states. J) Overlay of the intermediate (orange) and pre-PS (grey) map. Shown are two MTDs. The position of the adjacent B-tubule changes between the two maps. In the intermediate conformation, the next MTD (“n+1”) moves approximately 2nm away compared to the pre-state (indicated by black arrow). Furthermore the whole n+1 MTD rotates by around 6°.

In the pre-PS conformation, the dynein heads have moved ∼8nm proximally compared to the post-PS conformation (Figure 4D-F). In this pre-PS conformation, the linker is in a 90°-bent conformation, which is similar to the single particle cryo-EM analysis of cytoplasmic and axonemal dyneins heads in the presence of ADP-VO_4_ (Roberts et al., 2009; Schmidt et al., 2015), while the stalks are visible, pointing proximally towards the adjacent B-tubule (green and yellow in Figure 4G, respectively).

All these observations suggest that this is indeed the pre-PS conformation. From the subtomogram average as well as the surface rendered model one can see that the α and γ stalks are contacting to the adjacent B-tubule (Figure 5B,D,F,G,J).

**Figure 5.**
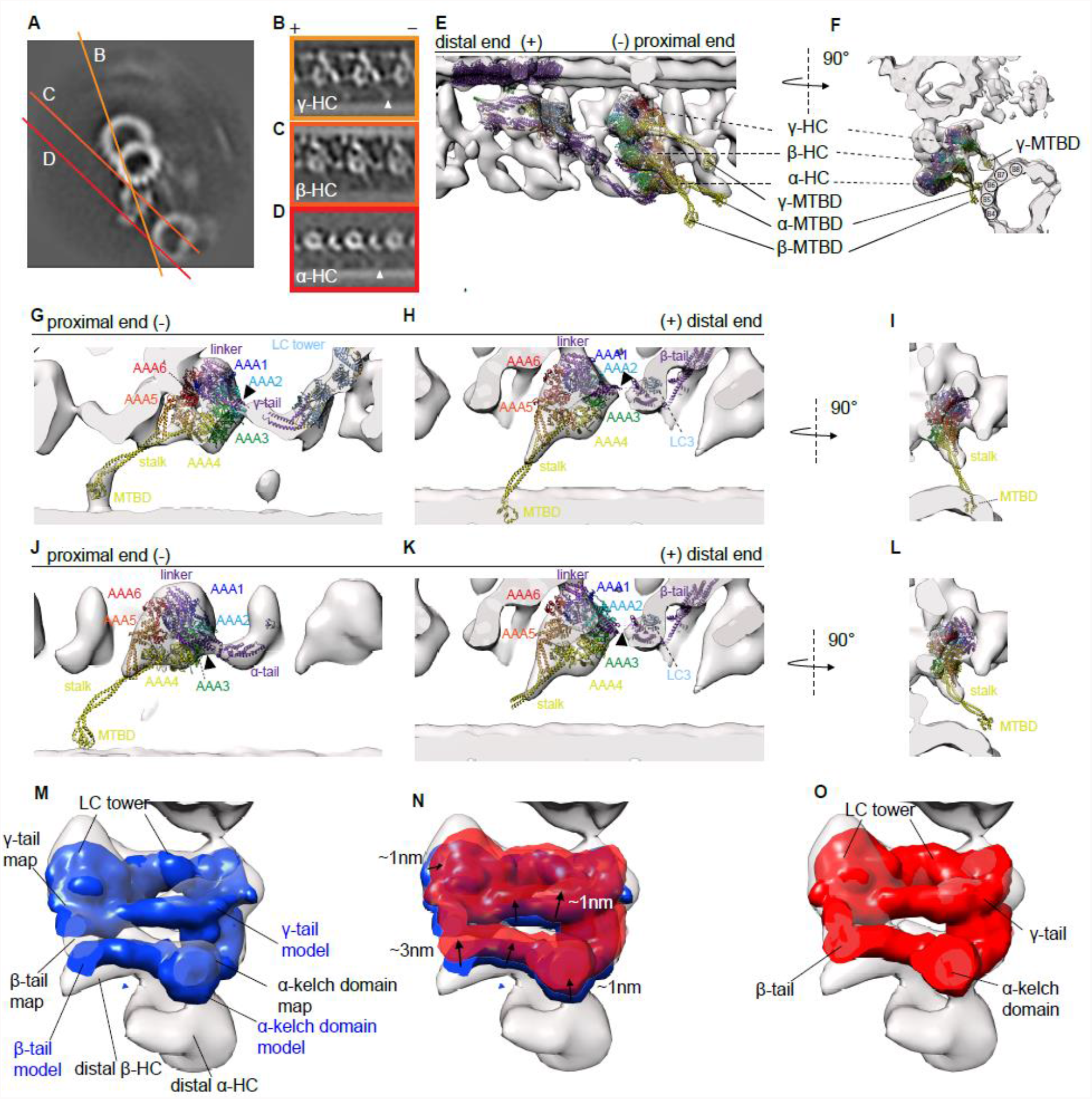
Pre-PS conformation of the ODA. (A) Cross section of one MTD in the pre-PS conformation. B-D) Tomographic slices for the γ, β and α HC respectively. The MTBD of the γ and α dynein is indicated with a white arrowhead. E-F) Two views of the overall structure of ODA in the pre-PS conformation with the pseudo atomic model fitted. (G-L) Enlarged view of individual dyneins in surface rendered representations with atomic models juxtaposed (PDB-4RH7 (G-I), PDB-6ZYW (L-K) and modified tail of PDB-7moq). The linker to tail connections are indicated by black arrowheads. The beginning of the stalk is visible for the β dynein (C, H-I, K-L). If the stalk had an extended conformation like in the cytoplasmic PDB-4RH7 conformation (Schmidt et al., 2015), the MTBD of the β HC would penetrates into the adjacent B-tubule (H-I). This is due to the dynein head position, which is too close to the adjacent B-tubule. The β stalk rather appears to be in a bend conformation which is much more similar to the conformation adapted in the Shulin-ODA complex (Mali et al., 2021) (K-L). M) 20Å resolution map of the post-PS tail complex (calculated from PDB-7MOQ) shown in a pre-PS tomographic map. N) 20Å resolution map of the pseudo-atomic pre-PS model overlaid on post-PS tail to highlight the changes. O) 20Å resolution map of the pseudo-atomic pre-PS model fitted in the pre-PS tomographic map.

The γ stalk of the pre-structure is in an extended conformation, attaching with its MTBD to the adjacent B-tubule between PF 7-8 (Figure 5B,F,G). The stalk of the β dynein appears to be in a curved conformation, similarly to the conformation it adapts in the Shulin-ODA complex (Mali et al., 2021) (Figure 5C, E,F, K-L). If this stalk is straight, the MTBD of β will be embedded into the B-tubule (Figure 5H,I). The MTBD of β is not clearly visible. Considering the conformation of the stalk it is most likely not bound to the adjacent B-tubule. A weak density between the PF5-6 on the adjacent B-tubule suggests that the MTBD of the α HC binds to the B-tubule with the α stalk in the unusual extended conformation (Figure 5D-F, J). With these stalk conformations the MTBD of the β dynein is not only more distally than the extended MTBD of the α dynein (Figure5E), but also more radially outwards (Figure 5F). In this case, the α and β MTBD would have to cross during a PS cycle.

With the combination of rigid body fitting of the crystallographically solved cytoplasmic dynein in the pre-PS and the recently solved dynein HC in the Shulin-ODA complex in which ATP binds, we were able to generate a pseudo-atomic model for the pre-PS state, which agrees with our tomographic maps (Figure 5F,K,L). The linker position of the cytoplasmic dynein coincides with the tomographic map of the pre-structure for the γ and β HC (Figure 4G-I). For these two HCs the linker conformation, spanning over AAA2, is the same as in the Shulin-ODA. The linker for the α dynein adopts a slightly different conformation from the cytoplasmic dynein. While the linker still has a 90° bend, it runs a little closer to AAA3, as seen in the Shulin-ODA complex. For our pre-PS map, the tail-linker connection can be seen for the γ and the α dynein head, but not for the β dynein (Figure 5G,H,J). The volume between the linker and the tail in the pre-PS and the intermediate structures corresponds to the number of missing residues. However, for the α dynein the empty volume between the linker and the kelch-like β propeller is too large to correspond only to the α linker. As already proposed in the previous section, this density could correspond to Lis1 or yet another still unidentified molecule. While in the post-PS this molecule has connection to the α tail and the AAA2/3, this unassigned density appears next to the Ig-Fln of the α tail, suggesting that during the PS cycle the connection to the dynein head is lost (Supplementary Figure 2Q).

While α- and β-HCs from Shulin-ODA, not cytoplasmic dynein, fit well to our density map, only γ-HC shows resemblance between cytoplasmic dynein and our map. The linker of the γ dynein goes over AAA2 and the stalk is in the straight conformation, which are both features of cytoplasmic dynein in the pre-structure (Figure 5G).

The tail complex of the post-PS ODA (PDB-7MOQ) fits overall to our pre-PS maps, meaning that the conformational change inside the tail complex is relatively small compared to the dynein head change.

The docking complex and the N-terminal region of the γ and β tail are very similar to the post-PS form in the pre-PS state. In the pre-PS conformation the proximal region from helical bundle 4 for γ tail and 5 for the β tail up to the dynein heads deviates from the post-PS conformation (Figure 5M). To fit to our cryo-ET map, the γ and β tail should move together with the LC-tower around 1nm proximally and 1nm closer to the A-tubule (Figure 5 N-O). The β tail at helical bundle 7-9 moves an additional 2nm closer to the γ tail (Figure 5 N). If this movement would not occur, the dynein head in the pre-PS conformation would sterically clash with the post-PS LC tower position. We summarized our modeling of the pre-PS in the supplementary movie 5.

Rao et al. have described the tail-to-head (TTH) interactions occurring in the post-PS of *Tetrahymena* ODA, and suggested that these TTH need to be broken for the PS to occur. We aimed to look at the intra- and inter-ODA interactions in the ODAs in the pre-PS conformation. The following densities connecting the dynein heads to other dynein heads within the same ODA (intra-ODA connection, indicated in Figure 6A-B yellow) and to the tail from the proximal ODA (inter-ODA connections, indicated in Figure 6A-B orange) were observed. In the pre-PS conformation the γ-HC density connects the LC tower via its AAA5/AAA6 site to LC8/LC10 (Figure 6B,C and D indicated by purple arrowhead). The stalk of the γ HC also comes very close to the LC tower at the region of LC8/6 (Figure 6D pink arrowhead). The intra-ODA density between the γ-HC and β-HC corresponds to the γ linker connecting to AAA2 of the β-HC (Figure 6A-C yellow arrowhead). There is a weak connection of the β-HC to the LC-tower in the region of LC8/10 (Figure 6C, indicated by a blue arrowhead). A more dominant connection of the C-terminal end of β linker is established to IC2 (Figure 6A-C, red arrowhead). In the case of the α-HC, the N-terminal region of the β linker is involved in connection to the AAA2/AAA3 region (Figure 6A,C green arrowhead). The α-HC shows a density connection to the proximal β-tail in the area of the kelch domain of the proximal α HC (Figure 6A-C, orange arrowhead).

**Figure 6.**
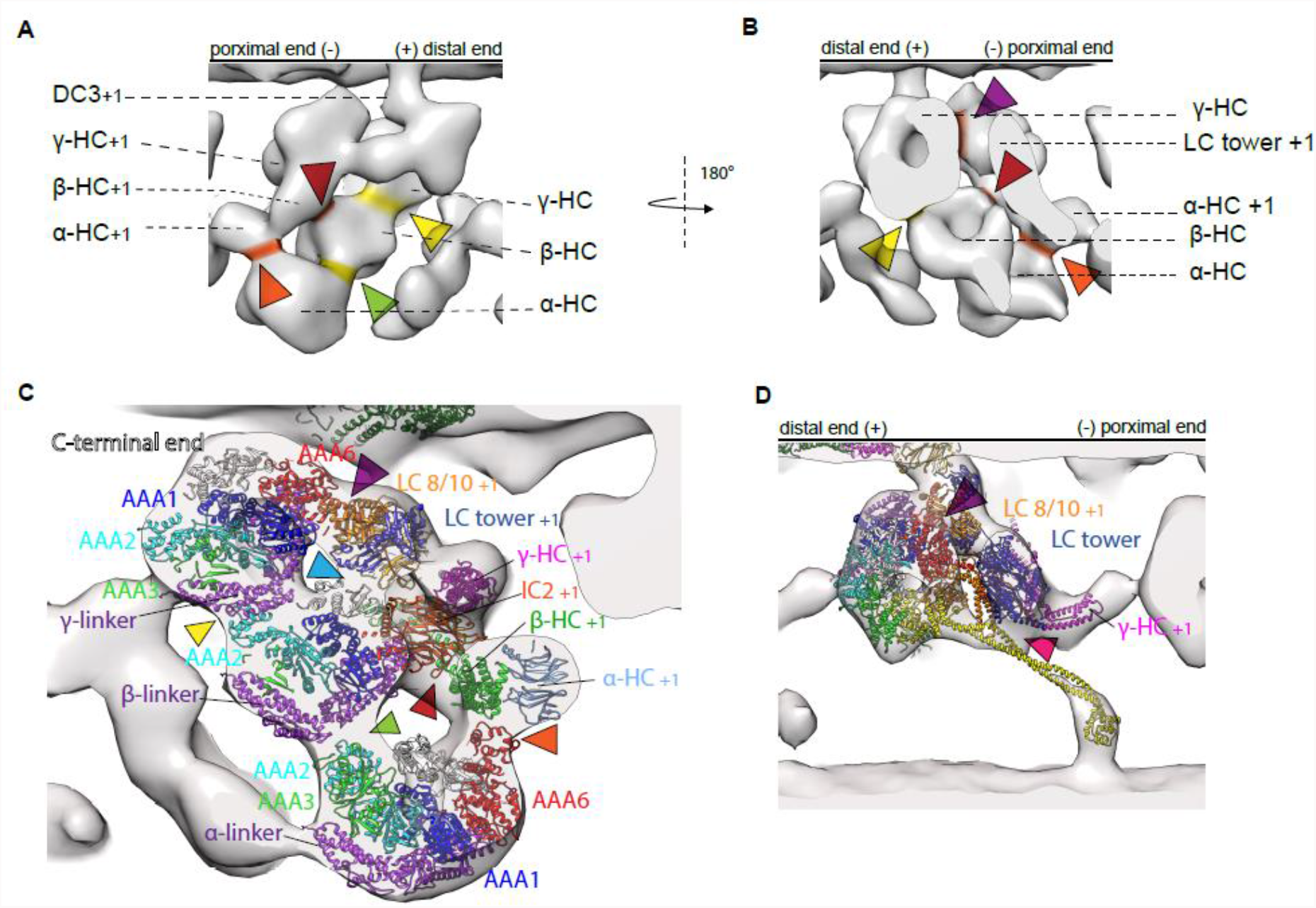
Intra and inter ODA connection in the pre-PS conformation. A-B) Surface rendered model of pre-PS conformation. Intra ODA connections between the three HCs are highlighted in yellow and inter ODA connections are highlighted in orange. C) Section through the three dynein chains. The pseudo-atomic structure of the pre-PS has been fitted into the cryo-ET map to highlight which parts of the ODA are involved in the interaction. The γ linker connects to the AAA2 from the β HC (yellow arrowhead). The β linker connects to the AAA2/AAA3 of the α HC (green arrowhead). AAA6 (purple arrowhead) and the stalk from the γ HC (pink arrowhead), as well as the C-terminus of the β HC (light blue arrowhead), interacts with the LC tower. The AAA6 from the β HC interacts with the proximal IC2 (red arrowhead). The AAA6 from the α HC interacts with the β tail of the proximal ODA (orange arrowhead). D) View of the γ HC with the PDBX-4RH7 highlighting the interaction between the γ HC and the proximal LC tower. AAA6 interacts at LC8/10 level (purple arrowhead) and the stalk interacts with LC6/8 (pink arrowhead).

The dynein heads of the intermediate structure are located between the post-PS and the pre-PS conformation (Figure 4E). The stalks of the dynein heads in this intermediate conformation are partially visible (Figure 7C-H). The base of the stalks, close to the dynein head are pointing proximally. Only for the γ dynein the entire stalk is weakly visible, including a density which corresponds to the MTBD of the γ dynein (Figure 7C). The γ stalk is in an extended conformation and the MTBD appears detached from the adjacent B-tubule. Although the linker of the intermediate conformation is not very evident, it appears that the linker of the γ and β HC are similar to the pre-PS linker conformation discussed above (Figure 4E,G,H). For the α HC the linker in the intermediate conformation seems to go over AAA3 (Figure 7H), something between the pre-like (AAA2/AAA3) and post-like (AAA3/AAA4) linker conformation. The unknown binding partner of the α HC is detached from the α head and bound to the α tail, as was seen for the pre-PS conformation (Supplementary Figure 2O and Q). Since the distance to the adjacent B-tubule is the same between the post-PS and the intermediate conformation, but the linker appears to have already changed to a pre-like conformation, the stalks can be in a straight conformation without protruding into the adjacent B-tubule (Supplementary Figure 2J-L). At the current resolution, the tail region resembles the post-PS conformation (Figure 7B). The TTH interactions in the intermediate conformation resemble the TTH interaction of the pre-PS conformation (Figure 7I,J). The γ dynein head detached from the NDD and forms now interaction with the LC tower of the proximally located ODA (Figure 7I,J orange indication). The γ linker interacts with the β head (Figure 7J, yellow arrowhead). Similar to the pre- and post-PS conformation the β dynein contacts the tail structure at IC2 (Figure 7I, J, light orange indication).

**Figure 7.**
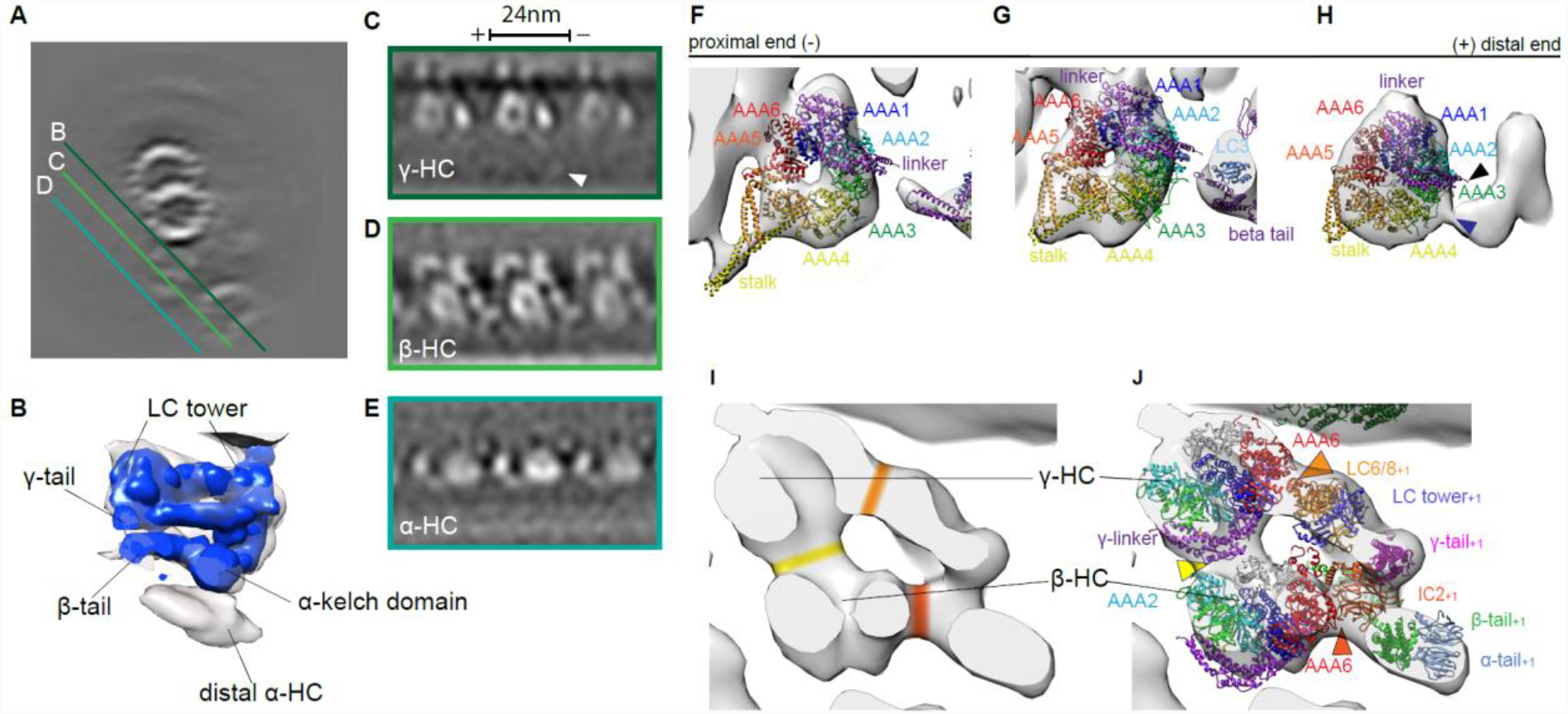
Intermediate conformation. A) Cross section of MTD. Indicated are the sections seen in C-E. C-E) Tomographic sections of γ, β and α HC respectively. (B) There are no major deviations between the tail structure of the post-model and the intermediate conformation tomographic map. F-H) Sections of the γ, β and α surface rendered map with the cytoplasmic dynein (PDB-4RH7) rigid body fitted. The tail complex from PBD-7MOQ is fitted as a rigid body. For α HC (H) the linker (purple) does adapt an intermediate conformation, whereas the linker spans over AAA3 (bottom blue arrow head) instead of going over AAA2 (top black arrow head). I) The intra ODA connection between the β head and the γ linker is highlighted in yellow. The inter ODA connections of the β dynein head to the IC2 and the connection of the γ head to the LC tower are indicated in orange. J) Atomic model fitted to the same view as in I, highlighting which proteins are involved in the TTH interactions. The highlighted regions from I are marked with arrows in J.

## Disscusion

In this study, we were able to show in situ structure of ODA in *Chlamydomonas* cilia by cryo-ET and demonstrated difference from in vitro structure in post-PS conformation by single-particle analysis. This difference could come from the preparation method. Where cryo-ET resembles the more native conformation and SPA has more purification steps involved, which can lead to higher resolution but also distort the native environment. The conformation of minimum energy when only one end is anchored on the A-tubule or binding to the B-tubule is different from the native conformation, where the ODA structure is restricted by both A- and B-tubules. Moreover, none of the SPA structures solved had all the components present. PDB-7KZM and PDB-7MOQ miss the adjacent B-tubule and PDB-7K5B and PDB-7K58 lack the docking complex and are assembled in vitro on MTDs. Removal of such opponent structures, which might exert some force on the ODA, could change the overall ODA structure. In the post-PS the MTBD is attached to the adjacent B-tubule. Without this binding partner, the stalk is highly flexible and can take conformations, which in the native environment it can most likely not obtain, such as the β stalk pointing distally found in PBD-7KZM. In contrast to all the SPA structures, we found all three HCs in a Post-2 conformation (according to the definition of (Kubo et al., 2021; Rao et al., 2021). Furthermore, an additional density appearing in the ODA in cryo-ET, which could be LIS1 bound to α-HC. Such smaller component can fall off during purification for SPA or smeared out during the averaging process due to flexibility.

Upon addition of ATP we found two major ODA conformation by unsupervised classification with RELION. One reason why no post-PS structure was present could be due to the higher nucleotide concentration compared to the previous work (Movassagh et al., 2010). Our intermediate conformation lacks the higher resolution information, such as the full-length stalk, which is why we speculate that the intermediate state represents a transitional state, where the linker already underwent the conformational change while the stalk is still finding the right PF to bind on the adjacent B-tubule. Our pre-PS map resembles the previously described ATP-VO_4_ state (Movassagh et al., 2010), we therefore assign this conformation to the ADP.Pi state, while the intermediate state probably represents the ATP state.

By cryo-ET analysis of cilia in the presence of ATP, we demonstrated how the interaction partner, B-tubule, influences the ODA confirmation. It becomes visible when we look at the different conformations during the PS cycle. The distance and angle of the neighboring MTD changed. This can be explained by the interaction of the adjacent B-tubule with the stalks and MTBDs. In the post-PS conformation, the stalks are upright and extended (Figure 2). This maintains a certain distance between two MTDs. Once ATP binds and subsequently the MTBD are released from the adjacent B-tubule all three dynein head move proximally by changing their linker position from post-to pre-PS conformation. The γ head breaks the TTH interaction with the NDD and forms new connections with the LC tower in the proximal ODA. During the whole PS cycle the interaction between the β HC and the IC2 seems to be kept. Although the resolution is not high enough to see changes inside IC2 directly, if assumed that IC2 does not shift with respect to the rest of the tail complex, it could serve as a rotation origin for the β-HC. The results in the intermediate conformation. To move from the intermediate to the pre-PS conformation, ATP is hydrolyzed and the MTBD bind the adjacent B-tubule. Differently from the model proposed by Lin et al. 2014 where the conformational change is done by rotation of the dynein head and stalk towards the distal end, our data suggests that the dynein heads move even more proximal than in the intermediate conformation, pushing the tail complex of the next ODA more proximal. By binding to the adjacent B-tubule the ODA can exert a force on the MTD which is bound by DC3, thereby pulling itself 2nm closer to the adjacent B-tubule. Pointing of the stalk of the β-dynein curved towards PF 4-5, (which was observed in previous findings in *sea urchin sperm* flagella by Lin et. al. 2014, Figure 3i), can introduce a torque, rotating the ODA relative to the adjacent MTD. Previously, from cytoplasmic dyneins, it is believed that the -PS happens in one go, this cannot be for the β dynein since the binding on PF seems to change between pre- and post structures. This opens up the question if the β dynein takes on a different role or there is another state after pre-PS and post-PS.

In all the three states and all the dyneins, the stalks extend to the proximal direction, which supports the winch hypothesis. However, the stalks are tilted more towards the proximal end in the pre-PS than the post-PS conformation, suggesting active rotation during the PS. The dynein heads change conformations in slightly different ways. The β dynein rotates around and moves with IC2 (Figure 4H, the connection to IC2 is indicated with black arrowheads) while γ and α do more translational movement (Figure 4G and I). While β rotates and shifts only very little compared to the other two HC, the α head can shift more proximally than β head, which could allow the α stalk to bind the PF more proximally than the β stalk (Figure 5E-F).

Recently it was found that the ODA are transported to the axoneme in an inactive Shulin-ODA complex. This structure was then solved to atomic resolution (Mali et al., 2021). Kubo et al. proposed how to open this Shulin-ODA complex to be in a post-PS conformation bound to the A-tubule. Our finding shows that the pre-PS conformation of the α and β-HCs in the Shulin-ODA complex and those from the ODA bound to the axonemes are almost the same. The β-HCs has similarity to the cytoplasmic dynein in the pre-PS state. With this we were able to build a pseudo-atomic pre-ODA. This model does not represent the precise atomic structure of the pre-PS ODA, but it can give important insight into the pre-PS conformation. This model shows that the linker in the axonemal pre-PS dynein is 90° and that the MTBD of γ and α bind on the adjacent B-tubule (Figure 5B, D, G-L). It furthermore highlights that a curved stalk conformation, as seen for the pre-PS β dynein, can natively occur. Apart from a global shift, and a local shift of the β tail closer to the γ tail in the pre state, our tomographic map was not resolved well enough to see structural changes in the tail complex. However, our data suggest that there is indeed some structural change happening in the tail complex during the PS cycle (Figure 5 M-O).

In conclusion, we have shown there are some differences between SPA and cryoET. Furthermore, we build a pseudo-atomic model of the pre-PS state (at intermediate resolution). We showed that the linker, head and stalks of the Shulin-ODA HC represent the native pre-PS ODA conformation.

## Materials and Methods

### Cells

*Chlamydomonas* cells (CC-125+) obtained from *Chlamydomonas* resource center were cultured in TAP medium, with a 12 hours dark and 12 hours light cycle. The cells were collected by centrifugation and deflagelated with dibucaine hydrochloride (Sigma-Aldrich, USA) following the Witman procedure (Witman, 1986). Flagella were collected by centrifugation at 22’000 g for 5 min. Axonemes were demembraned by 0.5% NP-40 treatment in 30 mM HEPES (pH 7), 5 mM MgSOs_4_, 1 mM DTT, 1 mM EGTA, 50 mM potassium acetate buffer. The activity of the purified axonemes was verified under the light microscope upon addition of 1mM ATP.

### EM grids

R3.5/1 Holey carbon copper grids (QUANTIFOIL, Germany) were glow-discharged by UV light. 10nm gold beads were applied to the grids as fiducial markers. Axoneme solution was mixed with ATP, to a final concentration of 1mM ATP, and immediately applied to the grids. These were manually backside blotted and plunge frozen on the Cryoplunge3 (Gatan, USA).

### Data collection

The tomographic data were acquired on a 300kV Titan Krios (Thermofisher, USA) at ETH Zürich with a K2 camera and GIF-Quantum energy filter (Gatan). Tilt series from -60° to 60° were collected with a 2° increment using a bidirectional tilt scheme starting from 0° for the data with ATP and unidirectional for the nucleotide free dataset using SerialEM (Mastronarde, 2005). The total electron dose used for both datasets was 60 electrons per Å^2^. The frames of the dose fractionated, normalized micrographs were aligned using IMOD alignframes. Tomograms were reconstructed using IMOD (Kremer et al., 1996). CTF correction was done in IMOD.

### Initial data processing – axoneme_aln

Subtomogram averaging was performed as described elsewhere (Bui and Ishikawa, 2013). This procedure includes manually picking of the MTDs in IMOD. The points along one MTD were interpolated and segmented into 24nm arrays. These subvolumes were aligned along one MTDs, assuming they have similar Euler angles. The 24nm subtomograms of all nine MTDs were then aligned based on the nine-fold pseudo symmetry of the axoneme. For obtaining the 96nm subunit, the right frame was selected from the 24nm segments, to reduce redundancy. More details can be found in Bui and Ishikawa (Bui and Ishikawa, 2013).

### Refined data processing – RELION

To eliminate reference bias, reference free classification and refinement in RELION (RELION version 2.1) was implied (Scheres, 2012). All 24nm subtomograms were extracted from the tomograms according to the aligned orientation and position determined by axoneme_aln, using BSOFT (Heymann, 2001). These prealigned subtomograms were imported into RELION (Workflow in Supplementary Figure 1A). There RELION 3D autorefine was used to refine the alignment of axoneme_aln. Then reference free 3D classification, without alignment, focused on the IDA and radial spokes and separately on one ODA of the 96nm repeat was carried out in RELION. A cubic mask generated with BSOFT included the three dynein heads and part of the tail complex for the ODA and another mask including all IDA and the radial spokes to separate the four different 96nm frames were applied. The outputs of the ODA classification were iteratively refined and re-classified with RELION and supervised classification (see below), resulting in three major classes. Further classification led to the classes being separated based on the missing wedge and therefore disregarded. The information of the 96nm frame and the ODA class was combined, to generate a 96nm subtomogram average for the different classes (Supplementary Figure 1B). Since none of these classes represented the post-PS conformation (apo), we used the data from a nucleotide free dataset and processed it in the same way as the data with ATP. The FSC was calculated in RELION using the same mask as was used for the last refinement step.

### Supervised classification

As a method for validation and further improvement of the class assignments, supervised classification (Movassagh et al., 2010) was implemented, the structures obtained from the reference free classification were used as references for the template matching based supervised classification. Only subtomograms with a cross-correlation value bigger than 0.15 were used for structure averaging.

### Data interpretation

The subaverages from the RELION jobs were visualized using IMOD and UCSF Chimera (Goddard et al., 2005). The recently published atomic models of the ODA structure by single particle cryo-EM (Walton et al., 2021, (PDB-7KZM) Kubo et al., 2021 (PDB-7MOQ) Rao et al., 2021 (PDB-7K58 and PDB-7K5B) were fitted into our 24nm subtomogram average and 96nm subtomogram average maps maximizing cross-correlation in UCSF Chimera (the list of the PDB files used in this study is in the supplementary table). The atomic models PBD-7MOQ, PDB-7KZM and PDB-7K58 were fitted as rigid bodies. The fit of the PDB-7KZM dynein heads was improved by individually rigid body fitting the dynein heads. For the PDB-7K5B, Coot (Emsley et al., 2010) was used for rigid body fitting of parts of the chains separately from the rest of the structure. Then real space refinement was used to reconnect the broken chains from the step before. For the pre-PS structures the crystallographic model of cytoplasmic dynein (PDB-4RH7) (Schmidt et al., 2015)and the HC of the Shulin-ODA complex (PDB-6ZYW) were fitted into the dynein head densities. With rigid body fitting and real space refinement the best fitting pre-PS dynein heads were used together with the PDB-7MOQ tail complex to build a pseudo-atomic model. For the visualization purpose in some figures, the function Segger in Chimera was used to hide B-tubule and further denoising in the intermediate state.

## Acknowledgements

We thank M. Peterek (ScopeM) and Daniel Boehlinger (CEMK) for technical assistance with electron microscopy, Prof. Masafumi Hirono (Hosei Univ., Japan) and Dr. Ken-Ichi Wakabayashi (Tokyo Inst. Technology) for advices about Chlamydomonas cell culture and axoneme isolation, Dr. Kai Zhang for providing the coordinate before publication and Alexandra E. Burger for constructive criticism of the manuscript. This work is supported by a grant from the Swiss National Science Foundation and Novartis Foundation for medical-biological research (NF310030_192644 and NF310030_192644). The authors declare no competing financial interests.

## Author contributions

AN and JM performed all the cell culturing and purification of axonemes. AN, JM and TI prepared cryo-ET samples and collected the cyro-ET datasets. NZ and JM processed the tomography data and NZ performed subtomogram averaging, classification and refinement. NZ built pseudoatomic models. NZ conceived the project. TI guided the project. NZ wrote the manuscript with help from all authors.

## Data and materials availability

Atomic coordinates of pseudo-atomic models and cryo-ET maps have been deposited in the Protein Data Bank under accession codes PDB-XXX (pseudo-atomic model post-PS),PDB-XXX (pseudo-atomic model pre-PS) and in the Electron Microscopy Data Bank under accession codes EMD-XXX,XXX,XXX.

## Supplementary Material

**Supplementary Figure 1.**
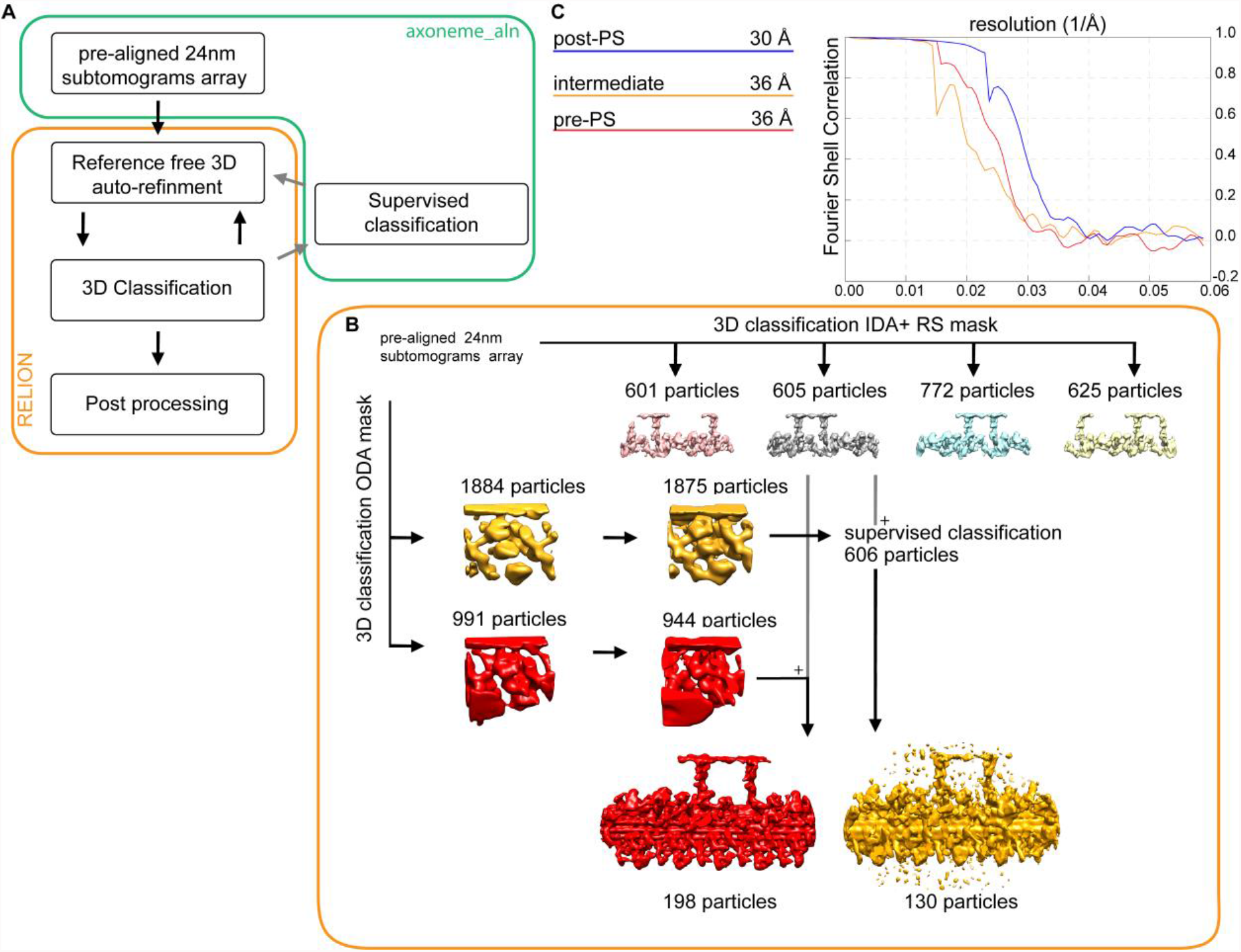
Workflow of subtomogram averaging. A) Pipeline of cryo-ET data processing. Two main software, axoneme_aln and RELION was used. B) Workflow in RELION with pre-aligned 24nm subtomograms as an input. Data processing was split into two parts, IDA and radial spoke classification and ODA classification. Recombination of these two information lead to 96nm subtomogram average. C) Post-processing of the 24nm subtomogram average lead to a FSC of 30Å for post-PS and 36Å for pre-PS and intermediate conformation.

**Supplementary Figure 2.**
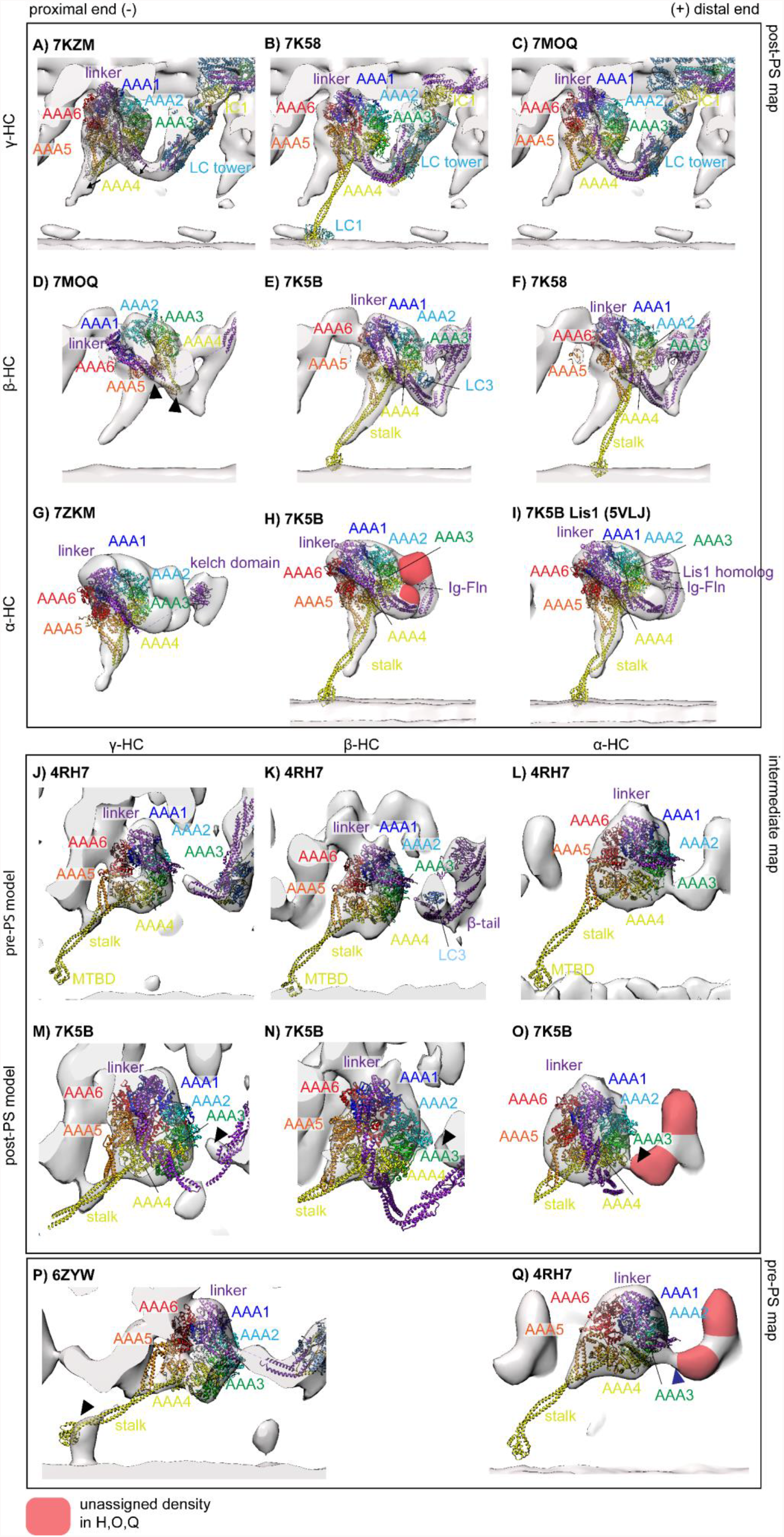
Suboptimal fitting of atomic resolution structures to our tomographic maps. A-C) Post-PS γ HC tomographic map with the atomic model of 7KZM, 7K58 and 7MOQ respectively. For A and C the stalk of the γ HC points to distally, due to the linker being in a Post-1 conformation for the atomic models. The arrows in A indicate the shift of fitting the dynein head separately from the tail complex. D-F) Post-PS β HC tomographic map with the atomic models of 7KZM, 7K58 and 7K5B. In D the linker and the stalk of the β dynein point into the same direction, indicated by black arrowheads. G-I) Post-PS α HC tomographic map with the atomic model of 7KZM and 7K5B. J-L) Intermediate tomographic map with the atomic model 4RH7 fitted to accommodate the stalk for the γ, β and α HC respectively. M-O) Intermediate tomographic map with the atomic model 7K5B fitted to accommodate the stalk for the γ, β and α HC respectively. Since the linker is in post-PS conformation, it does not align with the tomographic map, indicated by black arrowheads. P-Q) Pre-PS map of γ HC with the Shulin-ODA fitted as a rigid body. The head of the map and model of the γ HC are identical, but the stalk in the Shulin-ODA is bent, while the tomographic map suggest a straight conformation (black arrowhead). For the α HC, the cytoplasmic dynein was rigid-body fitted, the linker conformation of the model is slightly different from the one suggested by the tomographic map. The unassigned densities next to the α HC are highlighted in red for H,O and Q.

### Supplementary Movies

*Movie 1: Fitting 7KZM to our cryo-ET map, followed by the remodeled pseudo-atomic structure combining modified 7K5B and 7KZM*.

*Movie 2: Fitting 7MOQ to our cryo-ET map, followed by the remodeled pseudo-atomic structure combining modified 7K5B and 7KZM*.

*Movie 3: Two post-PS structures (7K58 and modified 7K5B) fitted to our post-PS cryo-ET structure*.

*Movie 4: Fitting 7K58 to our cryo-ET map, followed by the remodeled pseudo-atomic structure combining modified 7K5B and 7KZM*.

*Movie 5: Post-PS (modeling from modified 7K5B and 7KZM) and pre-PS structures, fitted to our cryo-ET maps*.

**Supplementary Table.** PDB data used for fitting and model building in this work.

|  |  |  |  |  |
| --- | --- | --- | --- | --- |
| 7KZM | Walton et al. | *Chlamydomonas* ODA anchored on A-tubule | No nucleotide | SPA |
| 7MOQ | Kubo et al. | *Tetrahymena* ODA anchored on A-tubule | No nucleotide | SPA |
| 7K5B | Zhang et al. | *Tetrahymena* ODA isolated and bound on B-tubule | No nucleotide | SPA |
| 7K58 | Zhang et al. | *Tetrahymena* ODA isolated and bound on B-tubule | No nucleotide | SPA |
| 6ZYW | Mali et al. | *Tetrahymena* ODA during IFT | ATP | SPA |
| 4RH7 | Schmidt et al. | *Homo sapiens* pre-power stroke dynein | ATP-VO<sub>4</sub> | crystallography |
| 5VLJ | Htet et al. | Yeast cytoplasmic dynein with *Homo sapiens* Lis1 | ATP-VO<sub>4</sub> | SPA |

